# Human cytomegalovirus glycoprotein variants governing viral tropism and syncytium formation in epithelial cells and macrophages

**DOI:** 10.1101/2024.02.12.580040

**Authors:** Giorgia Cimato, Xuan Zhou, Wolfram Brune, Giada Frascaroli

**Affiliations:** Leibniz Institute of Virology (LIV), Hamburg, Germany

**Author notes:** Address correspondence to Wolfram Brune.

## Abstract

Human cytomegalovirus (HCMV) displays a broad cell tropism, and the infection of biologically relevant cells such as epithelial, endothelial, and hematopoietic cells supports viral transmission, systemic spread, and pathogenesis in the human host. HCMV strains differ in their ability to infect and replicate in these cell types, but the genetic basis of these differences has remained incompletely understood. In this study, we investigated HCMV strain VR1814, which is highly infectious for epithelial cells and macrophages and induces cell-cell fusion in both cell types. A VR1814-derived bacterial artificial chromosome (BAC) clone, FIX-BAC, was generated many years ago but has fallen out of favor because of its modest infectivity. By sequence comparison and genetic engineering of FIX, we demonstrate that the high infectivity of VR1814 and its ability to induce syncytium formation in epithelial cells and macrophages depends on VR1814-specific variants of the envelope glycoproteins gB, UL128, and UL130. We also show that UL130-neutralizing antibodies inhibit syncytium formation, and a FIX-specific mutation in UL130 is responsible for its low infectivity by reducing the amount of the pentameric glycoprotein complex in viral particles. Moreover, we found that a VR1814-specific mutation in US28 further increases viral infectivity in macrophages, possibly by promoting lytic rather than latent infection of these cells. Our findings show that variants of gB and the pentameric complex are major determinants of infectivity and syncytium formation in epithelial cells and macrophages. Furthermore, the VR1814-adjusted FIX strains can serve as valuable tools to study HCMV infection of myeloid cells.

**Importance:** HCMV is a major cause of morbidity and mortality in transplant patients and the leading cause of congenital infections. HCMV infects various cell types, including epithelial cells and macrophages, and some strains induce the fusion of neighboring cells, leading to the formation of large multinucleated cells called syncytia. This process may limit the exposure of the virus to host immune factors and facilitate its spread. However, the reason why some HCMV strains exhibit a broader cell tropism and why some induce cell fusion more than others is not well understood. We compared two closely related HCMV strains and provided evidence that small differences in viral envelope glycoproteins can massively increase or decrease the virus infectivity and its ability to induce syncytium formation. The results of the study suggest that natural strain variations may influence HCMV infection and pathogenesis in humans.

## Introduction

Human cytomegalovirus (HCMV) is an opportunistic pathogen that is highly prevalent among human populations. In immunocompetent individuals, HCMV generally causes mild and self-limiting primary infections followed by lifelong persistence (1). However, it is a major cause of morbidity and mortality in immunocompromised individuals (2). HCMV is also the leading cause of congenital infection worldwide (3).

HCMV shows a broad cell and tissue tropism *in vivo* and can infect many organs and tissues in humans (4). *In vitro*, HCMV replicates efficiently in a variety of cell types, such as fibroblasts, endothelial, epithelial, and smooth muscle cells (5). Among the hematopoietic cells, HCMV has a tropism for cells of the myeloid lineage. Cells of the lymphoid lineage, by contrast, are not permissive. Myeloid cells play a crucial role in both lytic and latent infection (6–9). CD34+ hematopoietic progenitor cells and monocytes are an important reservoir for latent HCMV. Both cell types are not permissive for the lytic replication cycle. However, infected monocytes contribute to virus dissemination in the host by circulating in the bloodstream and migrating into tissues where they differentiate into macrophages or dendritic cells. Differentiation leads to HCMV reactivation from latency, production of infectious virus, and subsequent spread to other permissive cells (10–13).

HCMV cell tropism *in vitro* is governed by viral envelope glycoproteins that mediate receptor binding and virus entry into cells (14). Infection requires the fusion of the viral envelope with the host cell membrane. Key to this process are the glycoprotein B (gB) and the gH/gL complexes. Together, they form the core fusion machinery, which is highly conserved among the herpesviruses (15). gB is the primary fusogen and is responsible for membrane fusion. Its fusion activity is regulated by the gH/gL complex, which exists in two alternative forms in the virion envelope. The trimeric complex, consisting of gH, gL, and gO, interacts with platelet-derived growth factor alpha (PDGFRα) and mediates virus entry into fibroblasts (16, 17). Glycoproteins gH and gL can also form a complex with three glycoproteins encoded by the UL128 locus (UL128L). The pentameric complex (gH/gL/UL128/UL130/UL131A) interacts with the cellular receptors Neuropilin-2, OR14I1, or CD147 to allow virus entry into endothelial cells, epithelial cells, and macrophages (18–20). The relative amounts of the trimeric and pentameric complexes vary depending on the viral strain and the producer cell and affect the virus’ ability to infect specific cells and tissues (21). The core fusion machinery is also involved in cell-cell fusion and the formation of syncytia, which is induced to a variable degree by some HCMV strains (22).

AD169 and Towne are laboratory-adapted HCMV strains that have served for decades as workhorses of HCMV research (23). They have been passaged extensively in fibroblasts where they replicate to high titers. However, these strains have lost the tropism for endothelial cells, epithelial cells, and macrophages due to mutations in UL128L, resulting in the loss of the pentameric glycoprotein complex (24, 25). To preserve a broad cell tropism, scientists have propagated fresh clinical HCMV isolates in endothelial cells. Examples are the low-passage strains VHL/E, TB40/E, and VR1814 (26–28).

The cloning of HCMV genomes as bacterial artificial chromosomes (BAC) in *E. coli*, first reported in 1999 (29), has been a milestone in HCMV research. It allows the conservation of viral genotypes as BAC clones and greatly facilitates genetic engineering by BAC mutagenesis in *E. coli*. The first HCMV strains cloned as BACs were the laboratory strains AD169 and Towne (29–31). Shortly thereafter, the genomes of low-passage strains such as VR1814, TB40/E, TR, and Merlin were BAC-cloned as scientists recognized the need to study strains more closely reflecting the wild-type virus circulating in humans (32–35).

Strain VR1814 was isolated from a cervical swab of a pregnant woman with a primary HCMV infection (28). It was propagated in endothelial cells and has retained the ability to infect endothelial cells, epithelial cells, and macrophages with high efficiency. It has also retained the capability of being transferred from infected endothelial cells to polymorphonuclear leukocytes, which are thought to contribute to the hematogenous dissemination of HCMV (28). Moreover, VR1814 was shown to induce the formation of syncytia in infected epithelial cells (36). VR1814 was the first low-passage HCMV strain to be cloned as a BAC, and the VR1814-derived BAC clone was named FIX-BAC (32). While FIX (the virus reconstituted from FIX-BAC) was able to infect endothelial and myeloid cells, its infectivity for these cell types was low, and it did not produce high titers of cell-free virus. Hence, FIX-BAC did not gain wide acceptance in the field, and BACs of other low-passage strains were used more frequently.

In the present study, we investigated the genetic basis of the pronounced tropism of VR1814 for epithelial cells and macrophages. By sequence comparison and genetic engineering of FIX, we demonstrated that specific variants of gB and UL128L are responsible for the high infectivity of VR1814 and its ability to induce cell-cell fusion. Moreover, we showed that a VR1814-specific mutation in US28, encoding a membrane protein important for latency maintenance, further increases the virus’ ability to infect macrophages.

## Results

### HCMV strain VR1814 is highly infectious for several cell types and induces cell-cell fusion

HCMV strain VR1814 has been used in numerous studies because of its broad tropism for various cell types, including macrophages (28, 32, 36, 37). To quantify its infectivity, we infected different cell types with VR1814 and TB40/E, another widely used clinical strain. We also compared VR1814 with FIX, a virus reconstituted from FIX-GFP BAC (38). Two days after infection, cells were analyzed by immunofluorescence for the presence of the viral immediate-early proteins IE1 and IE2. Strains VR1814 and TB40/E infected similar percentages of fibroblasts, epithelial, and endothelial cells (Fig. 1A-C). However, VR1814 showed a higher infection rate than TB40/E in THP-1-derived macrophages and monocyte-derived M1 macrophages (Fig. 1D, E), but a similar infectivity for monocyte-derived M2 macrophages (Fig. 1F). In contrast, FIX showed very low infection rates for all cell types besides fibroblasts (Fig 1A-F).

**Figure 1.**
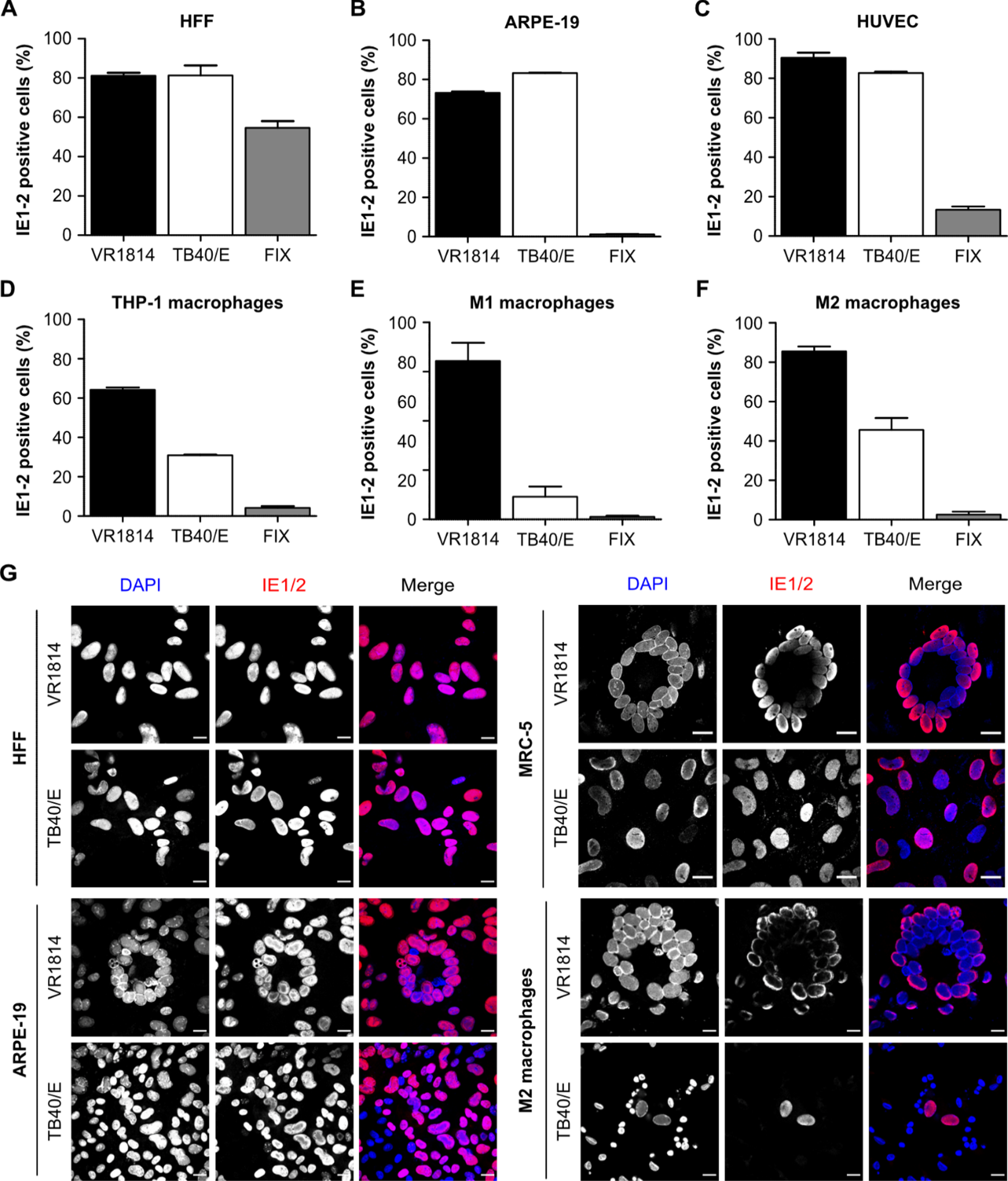
HCMV infectivity and syncytium formation in different cell types. HFF and ARPE-19 cells were infected with HCMV strains VR1814, TB40/E, and FIX at an MOI of 1. HUVEC cells, THP-1-derived macrophages, and M1 and M2 macrophages were infected at an MOI of 5. (A to F). The percentage of infected cells was determined by immunostaining with an antibody recognizing IE1 and IE2. Bars represent the mean ± SEM of three independent experiments. (G) Syncytium formation was analyzed in the indicated cell types at 5 dpi. Cell nuclei were counterstained with DAPI. Scale bar, 20 µm.

VR1814 was reported to induce cell-cell fusion in infected MRC-5 fibroblasts and ARPE-19 epithelial cells, leading to the appearance of large multinucleated cells. These syncytia usually display a circular arrangement of nuclei akin to the petals of a flower (36, 39, 40). To compare the abilities of VR1814 and TB40/E to induce cell-cell fusion, we infected fibroblasts, epithelial cells, and macrophages and analyzed syncytium formation on day 5 post-infection by immunofluorescent staining for IE1/2. In epithelial cells and macrophages, large syncytia were observed upon VR1814 infection (Fig. 1G). Interestingly, fibroblasts differed in their susceptibility to VR1814-induced cell-cell fusion. Numerous large syncytia were observed in MRC-5 fibroblasts, consistent with previous reports (39, 40), but not in HFF (Fig. 1G). This result suggested that HFF fibroblasts are less susceptible to HCMV infection-induced cell-cell fusion. Similarly, a previous study has found HFF to be less susceptible than MRC-5 cells to fusion induced by overexpression of viral envelope glycoproteins (41). In contrast to VR1814, TB40/E induced little or no syncytium formation in any of the cell types analyzed (Fig. 1G).

### Genetic differences between VR1814 and FIX

FIX-BAC was obtained by cloning the genome of VR1814 as a BAC in *E. coli* (32). However, as shown in Figure 1, FIX has a much lower infectivity than VR1814 for most cell types. It also does not induce syncytium formation in MRC-5 cells as VR1814 does (39). A single nucleotide polymorphism in the gB gene, leading to a serine-to-glycine exchange at amino acid 585 (S585G), was shown to be responsible for the increased fusogenicity of VR1814 gB (39). However, the reason for the relatively poor infectivity of FIX in epithelial cells, endothelial cells, and macrophages has remained unknown.

To identify the genetic basis of these differences, we determined the genome sequences of VR1814 and the FIX-GFP, a derivative of FIX-BAC containing a GFP expression cassette within the BAC vector backbone (38). Although the genome sequences of both viruses were published several years ago (34, 42), we wanted to compare the genomes of our isolates. Therefore, we determined the complete sequences of the FIX-GFP BAC and our VR1814 virus (hereafter called VR1814_GF_) by Illumina sequencing. As expected, the viral sequences of FIX-GFP and FIX-BAC were identical. The VR1814_GF_ sequence was most similar to the sequence of an endothelial cell-adapted VR1814 sequenced by Dargan et al. (42). Compared to FIX, the VR1814_GF_ sequence showed several differences apart from the known deletion of genes IRS1 to US6 in FIX, which was introduced during BAC cloning (32, 34). A summary of the differences in coding regions of the genome is listed in Table 1.

**Table 1.** Genetic differences in coding regions of HCMV VR1814_GF_ compared to FIX.

| gene | protein | location* | sequence<sup>†</sup> | coding effect |
| --- | --- | --- | --- | --- |
| UL8 | UL8 | 164499 | A | Substitution (E212K) |
| UL25 | UL25 | 32046 | CC+ | Frameshift |
| UL30 | UL30 | 37446 | A+ | Frameshift |
| UL31 | UL31 | 38319 | C | Substitution (V44A) |
| UL38 | UL38 | 51477 | GGG | In-frame insertion |
| UL41A | UL41A | 54327 | T | Substitution (E58K) |
| UL43 | UL43 | 55889 | C | Substitution (T143A) |
| UL50 | NEC2 | 73746 | GAG | In-frame insertion |
| UL55 | gB | 82890 | C | Substitution (S585G) |
| UL56 | TRM1 | 85615 | T | Substitution (V515M) |
| UL57 | DNBI | 89021 | C | Substitution (N803S) |
| UL75 | gH | 111133 | G | Substitution (L43P) |
| UL76 | UL76 | 111993 | C | Substitution (L180P) |
| UL80.5 | APNG | 117852 | CGC | In-frame insertion |
| UL89 | TRM3 | 133963 | G | Substitution (C547R) |
| UL95 | UL95 | 140692 | GGT+ | In-frame insertion |
| UL98 | UL98 | 144900 | G | Substitution (K352E) |
| UL98 | UL98 | 145063 | C | Substitution (D406A) |
| UL100 | gM | 147480 | C | Substitution (P3R) |
| UL104 | UL104 | 151475 | G | Substitution (L590P) |
| UL111A | vIL-10 | 160884 | T+ | Frameshift |
| UL111A | vIL-10 | 160925 | GAC | In-frame insertion |

| gene | protein | location* | sequence† | coding effect |
| --- | --- | --- | --- | --- |
| UL112 | UL112 | 163725 | GGT | In-frame insertion |
| UL116 | UL116 | 166522 | GGC | In-frame insertion |
| UL121 | UL121 | 170032 | C+ | Frameshift |
| UL122 | IE2 | 171135 | A | Substitution (S376F) |
| UL122 | IE2 | 171910 | AGG | In-frame insertion |
| UL128 | UL128 | 176683 | C | Substitution (F33V) |
| UL130 | UL130 | 177250 | G | Substitution (P72S) |
| UL130 | UL130 | 176856 | T | Substitution (T203N) |
| UL147A | UL147A | 180059 | C | Substitution (Y55X) |
| UL147 | vCXCL2 | 180375 | A+ | Frameshift |
| UL135 | UL135 | 188726 | G | Substitution (W284R) |
| UL135 | UL135 | 189267 | CTA | In-frame insertion |
| UL133 | UL133 | 189770 | GTC | In-frame insertion |
| UL150A | UL150A | 193227 | GGC | In-frame insertion |
| US15 | US15 | 209669 | G | Substitution (F87L) |
| US20 | US20 | 214013 | T | Substitution (W231X) |
| US28 | US28 | 225076 | GAC | In-frame insertion |
| US28 | US28 | 225998 | TT+ | Frameshift |
| US33A | US33A | 230110 | C | Substitution (L39P) |
| US34 | US34 | 230525 | GA+ | Frameshift |
| TRS1 | TRS1 | 234246 | G | Substitution (E35A) |

\* Nucleotide position in the genome sequence of VR1814 (GenBank GU179289).
† The mutated nucleotide at each location is shown.

### VR1814-specific gB and UL128L variants increase the infectivity of FIX in epithelial cells

We wanted to find out which of the sequence variations (Table 1) were primarily responsible for the higher infectivity of VR1814 in epithelial cells and macrophages. As the differences were too many to test them all, we decided to focus on a few that appeared to be strong candidates:

i. The S585G substitution in gB, which was previously shown to increase the fusogenicity in MRC-5 fibroblasts (39, 40).
ii. The three substitutions in UL128L (F33V in UL128, P72S, and T203N in UL130), as the pentameric glycoprotein complex is required for infection of epithelial cells and macrophages.
iii. The frameshift mutation in US28, leading to a short missense amino acid sequence after aa 314, as a previous paper has described an altered signaling activity of a truncated US28(1–314) protein (43).

To test which of these gene variants contribute to the phenotypic differences between VR1814_GF_ and FIX, we generated a set of FIX mutants by BAC mutagenesis. They carry VR1814_GF_ -specific variants of UL55/gB (“B”), of the pentamer components UL128 and UL130 (“P”), and the chemokine receptor US28 (“C”). Three single, two double, and one triple mutant were constructed (Fig. 2A).

**Figure 2.**
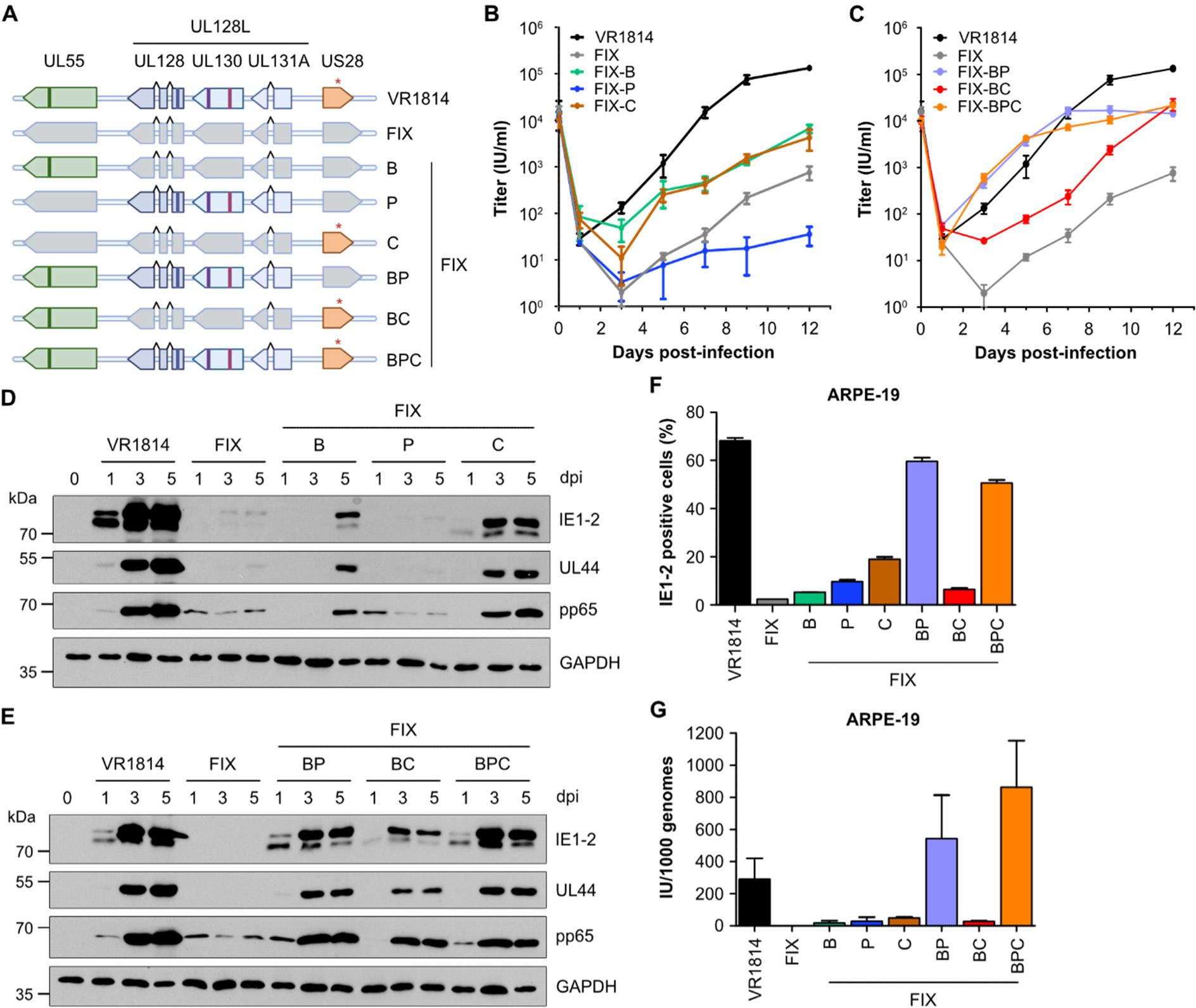
Replication and infectivity of recombinant FIX strains on ARPE-19 cells. (A) Schematic of the recombinant FIX strains generated by BAC mutagenesis. They contain VR1814_GF_-specific variants of UL55/g<u>B</u>, the <u>P</u>entamer proteins UL128 and UL130, and the US28 <u>C</u>hemokine receptor. (B, C) ARPE-19 cells were infected at an MOI of 0.5. Supernatants were harvested at different times post-infection and titered on HFF cells. Viral titers are shown as mean ± SEM of three biological replicates. (D, E) ARPE-19 cells were infected at an MOI of 1. Whole-cell lysates were harvested 1, 3, and 5 dpi, and the levels of the viral proteins IE1 and IE2, UL44, and pp65 were analyzed by immunoblotting. (F) ARPE-19 cells were infected at an MOI of 1. Relative infectivities of the recombinant FIX strains were determined at 1 dpi as percentage of IE1/IE2-positive nuclei. Mean ± SEM of three independent experiments. (G) The infectivities of the virus stocks on ARPE-19 cells were determined by quantifying viral genome copies by qPCR and infectious units (IU) by titration. Infectivity is shown as infectious units (IU) per 1000 viral genomes. Mean values ± SEM are shown.

First, we tested the ability of the recombinant FIX strains to replicate and spread in ARPE-19 epithelial cells. In multistep replication kinetics, the single mutants FIX-B and FIX-C replicated to higher titers than the parental FIX strain (Fig. 2B). Surprisingly, FIX-P replicated to lower titers than FIX. Among the double and triple mutants, FIX-BP and FIX-BPC replicated most efficiently in ARPE-19 cells. Up to day 7 post-infection, their titers were at least as high as those of VR1814_GF_, although the peak titers at 9 and 12 dpi were approximately 5 to 9-fold lower than those of VR1814_GF_ (Fig. 2C). We also determined viral protein expression on days 1, 3, and 5 post-infection (Fig. 2D, E). The results were consistent with those of the growth curves.

Next, we determined the relative infectivity of the recombinant viruses on ARPE-19 cells. The cells were infected at an MOI of 1 (based on titers determined on HFF cells), and the percentage of IE1/2-positive cells was determined at 1 dpi by immunocytochemistry (Fig. 2F). Again, the FIX-BP and FIX-BPC mutants had the highest relative infectivity, almost reaching the infectivity of VR1814_GF_. We also determined the absolute infectivity on ARPE-19 cells of each viral stock as infectious units (IU) per 1000 genomes. The results showed that all recombinant FIX strains had a higher infectivity than the parental virus and that VR1814_GF_, FIX-BP, and FIX-BPC had the highest infectivities (Fig. 2G).

### VR1814-specific variants of gB and UL128L promote cell-cell fusion and syncytium formation in epithelial cells

To assess the ability of the recombinant FIX strains to induce cell-cell fusion and syncytium formation in epithelial cells, ARPE-19 cells were infected at an MOI of 1. At 5 dpi, cells were fixed and immunostained for IE1/IE2. Syncytium formation was analyzed by microscopy (Fig. 3A). The results showed that only VR1814_GF_, FIX-BP, and FIX-BPC were capable of inducing cell-cell fusion and the formation of large multinucleated syncytia. To quantify syncytium formation, we used a previously established system based on a dual split protein (DSP) consisting of split GFP and split *Renilla* luciferase (40). ARPE-19 cells stably expressing either DSP1-7 or DSP8-11 were mixed (1:1 ratio), seeded in 96-well plates, and infected with HCMV strains. Cell-cell fusion was quantified by measuring *Renilla* luciferase activity in cells at 5 dpi. High luciferase activities were detected only in cells infected with VR1814_GF_, FIX-BP, and FIX-BPC (Fig. 3B), consistent with our observations by microscopy (Fig. 3A). These results suggested that syncytium formation in ARPE-19 cells requires VR1814_GF_-specific variants of both gB and UL128L.

**Figure 3.**
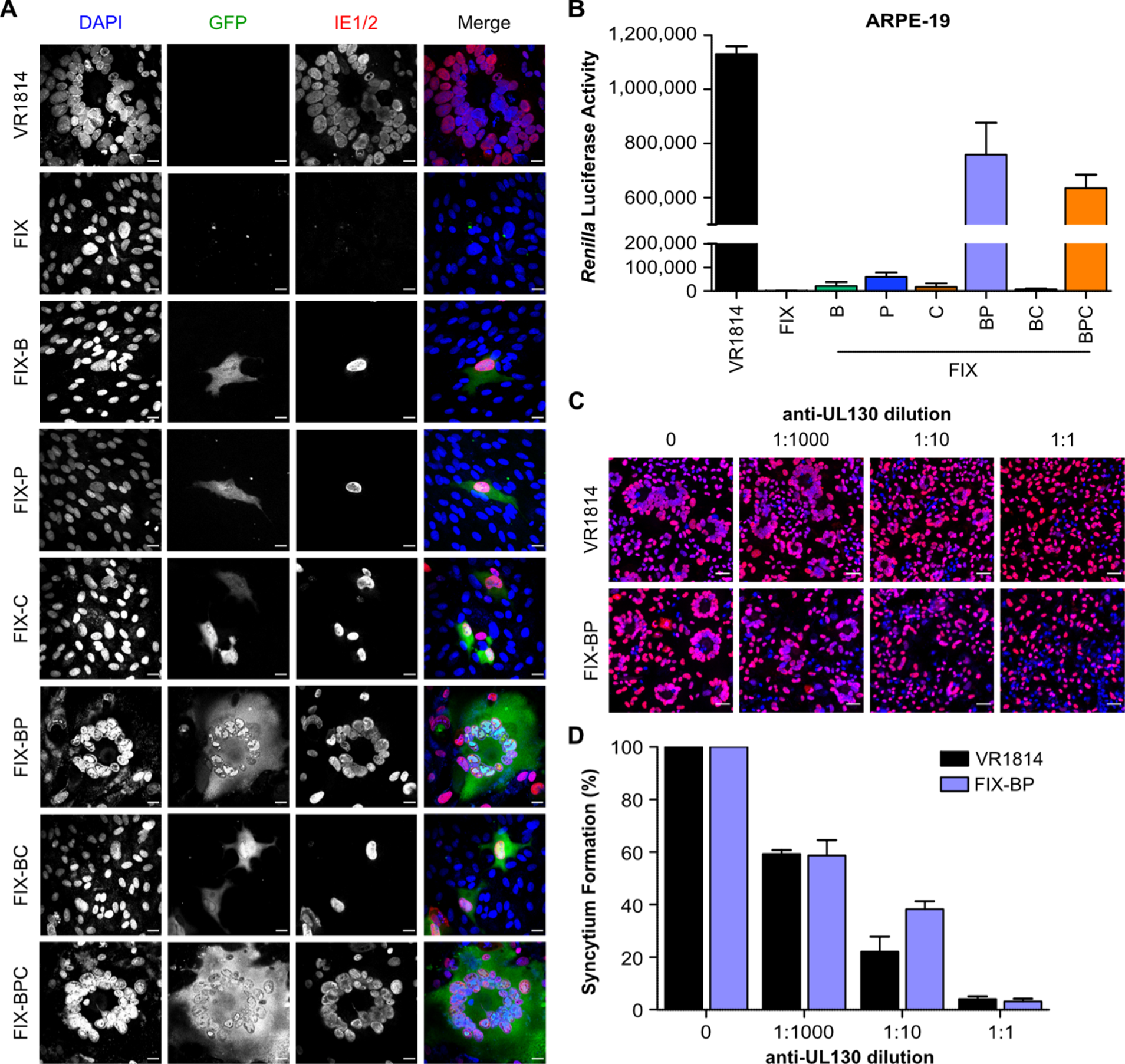
Epithelial cell fusion induced by VR1814_GF_ and recombinant FIX strains. (A) ARPE-19 cells were infected at an MOI of 1, fixed at 5 dpi, and stained with an anti-IE1/2 antibody (red). Nuclei were counterstained with DAPI (blue). Syncytium formation was analyzed by microscopic inspection. All FIX strains express GFP. Scale bar, 20 μm. (B) DSP-expressing ARPE-19 cells were infected as described above. *Renilla* luciferase activity was measured at 5 dpi. Mean ± SEM of three independent experiments. (C, D) DSP-expressing ARPE-19 cells were infected with VR1814_GF_ or FIX-BP and incubated with dilutions of an anti-UL130 hybridoma supernatant. (C) Syncytium formation was observed microscopically on day 5 post-infection and (D) quantified by measuring *Renilla* luciferase activity. Mean ± SEM of three independent experiments.

To further confirm the requirement of the pentameric complex for cell-cell fusion, we infected DSP-expressing ARPE-19 cells with VR1814_GF_ and FIX-BP as described above and treated the cells with a neutralizing anti-UL130 monoclonal antibody (44). Three hours post-infection, the virus inoculum was removed and replaced with medium containing serial dilutions of an anti-UL130 hybridoma (clone 3E3) supernatant. At 5 dpi, the formation of very large syncytia was observed only in the untreated infected cells. The size and number of syncytia decreased with increasing amounts of hybridoma supernatant (Fig. 3C). For a more objective quantification of syncytium formation, we measured *Renilla* luciferase activity in cells and calculated the relative inhibition by the anti-UL130 antibody (Fig. 3D). Increasing amounts of the monoclonal antibody inhibited cell-cell fusion by either virus. The results confirmed the crucial role of UL128L in HCMV-induced epithelial cell fusion.

### FIX-derived variants of UL128 and UL130 impair infectivity and cell-cell fusion in ARPE-19 cells

FIX-BP contains three VR1814_GF_-specific amino acid exchanges in UL128 (F33V) and UL130 (P72S and T203N) (Fig. 2A and Table 1). To find out to which extent these amino acid exchanges in UL128L contribute to viral infectivity and virus-induced cell-cell fusion, we reverted each of the three positions in FIX-BP individually back to the FIX variant by BAC mutagenesis (Fig. 4A). The three recombinant strains FIX-BP(V33F), FIX-BP(S72P), and FIX-BP(N203T) were first analyzed on ARPE-19 cells in terms of relative infectivity. As shown in Figure 4B, each of the three revertant FIX strains had lower infectivity on epithelial cells compared to the parental FIX-BP strain. However, the strongest reduction in infectivity was observed with FIX-BP(S72P), whose infectivity was very similar to FIX (Fig. 4B). Next, we measured cell-cell fusion induced by these viruses by using DSP-expressing ARPE-19 cells. Again, the ability of FIX-BP to induce cell-cell fusion was abolished by the S72P exchange in UL130 (Fig. 4C, D).

**Figure 4.**
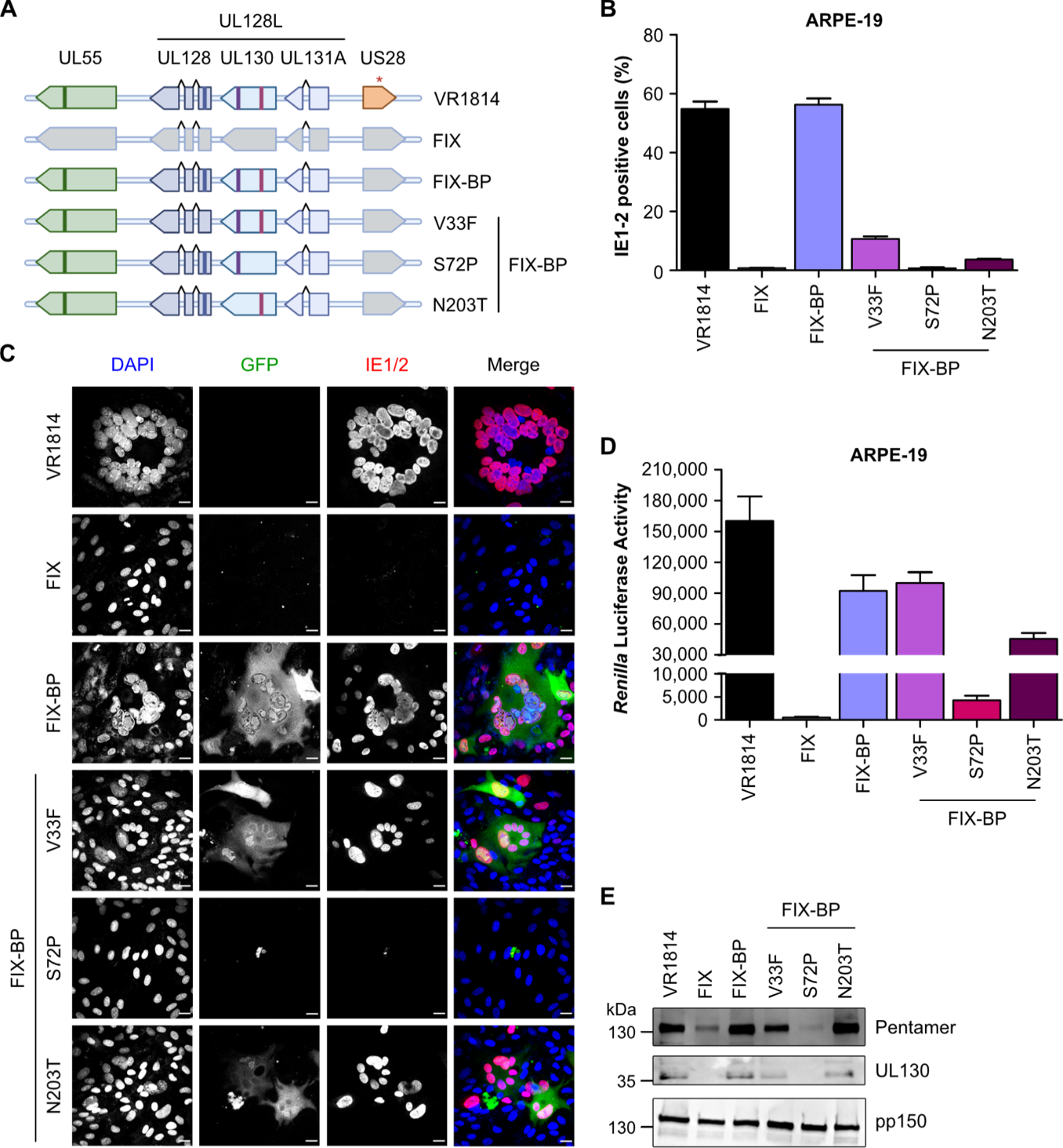
Impact of VR1814_GF_ and FIX-specific variants of UL128L on infectivity and cell-cell fusion. (A) Schematic of the recombinant FIX strains. (B) ARPE-19 cells were infected at an MOI of 1. Relative infectivities of the recombinant FIX strains were determined at 1 dpi as percentage of IE1/IE2-positive nuclei. Mean ± SEM of three independent experiments. (C, D) Syncytium formation in DSP-expressing ARPE-19 cells was observed at 5 dpi. (C) Cells were stained with an anti-IE1/2 antibody and nuclei were counterstained with DAPI. Scale bar, 20 μm. (D) Syncytium formation was quantified by measuring *Renilla* luciferase activity. Mean ± SEM of three independent experiments. (E) Lysates of purified virions were analyzed by immunoblot. The pentameric complex, separated on a non-denaturing gel, was detected with an anti-UL128 antibody. UL130 and pp150 (loading control) were separated on a denaturing gel and detected with specific antibodies.

Amino acid exchanges in viral envelope glycoproteins may affect infectivity and fusogenicity either by changing their incorporation in viral particles or their interaction with cellular receptors. To measure the incorporation of the pentameric glycoprotein complex into viral particles, we analyzed purified virus stocks by immunoblot analysis. The pentameric complex was detected by electrophoretic separation in a non-denaturing polyacrylamide gel and immunoblot analysis with a UL128-specific antibody. The UL130 protein was detected by standard SDS-PAGE and a UL130-specific antibody (Fig. 4E). Input was normalized to the capsid-associated tegument protein pp150. We also attempted to use the major capsid protein (MCP) for normalizing the input. However, this did not work as the available anti-MCP antibody reacted very poorly with MCP of VR1814 and FIX. The results showed very low quantities of the pentameric complex in purified virus preparations of FIX and FIX-BP(S72P) (Fig. 4E). This is consistent with a previous report by Murrell et al., which has shown low pentamer levels in FIX and a recombinant HCMV Merlin strain carrying UL128L of FIX (45).

### VR1814_GF_-specific variants of gB and UL128L increase infectivity and fusogenicity in macrophages

To assess the ability of the recombinant FIX viruses to infect macrophages, we first used THP-1-derived macrophages, an established model for HCMV infection of this cell type (46–48). THP-1 monocytes were treated with phorbol 12-myristate 13-acetate (PMA) for 3 days to induce their differentiation into macrophages. THP-1-derived macrophages were infected at an MOI of 5 with VR1814_GF_ and recombinant FIX strains. On days 1 and 5 post-infection, cells were fixed, and the percentages of IE1/2-expressing cells were determined by immunofluorescent staining. VR1814_GF_, FIX-BP, and FIX-BPC infected macrophages much more efficiently than FIX and the FIX single mutants (Fig. 5A, B), similar to what was previously observed in ARPE-19 epithelial cells (Fig. 2). Interestingly, FIX-BC infected THP-1-derived macrophages with similar efficiency as FIX-BP and FIX-BPC, contrary to what was previously observed in epithelial cells. This result suggested that the C-terminally truncated US28 protein promotes infection and IE gene expression in THP-1-derived macrophages.

**Figure 5.**
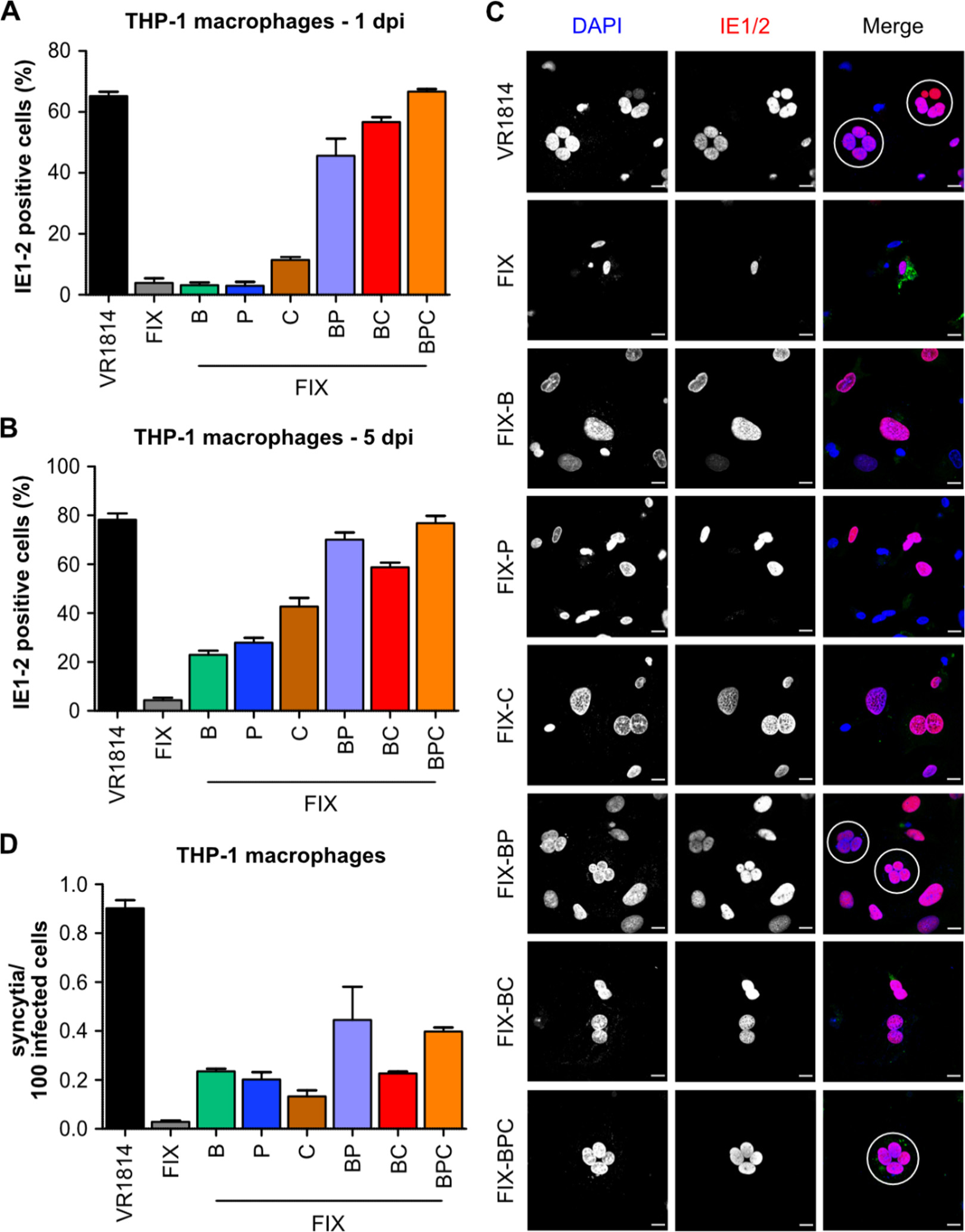
Infectivity of recombinant FIX strains and ability to induce syncytium formation in THP-1-derived macrophages. (A, B) THP-1 cells were differentiated into macrophages by PMA treatment for 3 days and infected at a MOI of 5. Relative infectivities of the recombinant FIX strains were determined at 1 and 5 dpi as percentage of IE1/2-positive nuclei. Mean ± SEM of three independent experiments. (C) THP-1-derived macrophages were infected at an MOI of 5, and syncytium formation was analyzed on day 5 post-infection. Cells were stained with an anti-IE1/2 antibody and nuclei were counterstained with DAPI. Syncytia are marked by white circles. Scale bar, 20 μm. (D) Syncytium formation was quantified at 5 dpi by microscopic inspection and counting. Mean ± SEM of three independent experiments.

Since VR1814_GF_ can induce syncytium formation in macrophages (Fig. 1G), we tested the ability of the recombinant FIX strains to do the same. THP-1-derived macrophages were infected at an MOI of 5, and syncytium formation was visualized microscopically (Fig. 5C). Only THP-1-derived macrophages infected with VR1814_GF_, FIX-BP, and FIX-BPC showed syncytium formation at 5 dpi. Unfortunately, we could not use the DSP system in THP-1-derived macrophages as it did not work reliably in this cell type. Apparently, the expression of the DSP halves did not remain stable upon passaging and differentiation of the transduced THP-1 cells. Therefore, syncytium formation had to be evaluated by microscopic inspection and counting. As shown in Fig. 5D, syncytium formation was induced most vigorously by VR1814_GF_, FIX-BP, and FIX-BPC, suggesting that VR1814_GF_-specific variants of gB and UL128L are needed for the fusion of macrophages.

Even though THP-1-derived macrophages are an established and widely used model for primary human macrophages, we wanted to verify our observations with macrophages derived from human peripheral blood monocytes. The monocytes were differentiated into M1 or M2 macrophages and infected in the same way as the THP-1-derived macrophages. Due to limited material, we did not include the FIX single mutants FIX-B, FIX-P, and FIX-C as none of them had a striking phenotype in the previous experiments. Similar to the results with THP-1-derived macrophages (Fig. 5A), FIX-BP and FIX-BPC mutants had the highest relative infectivities on both M1 and M2 macrophages, almost reaching VR1814_GF_ levels (Fig. 6A, B). Interestingly, we observed increased infectivity of FIX-BC in M2 macrophages, but not in M1 macrophages, suggesting a possible role of the truncated US28 protein in HCMV infection of macrophages depending on their polarization state. We also investigated syncytium formation at 5 dpi by microscopic inspection and counting. Consistent with previous observations in THP-1-derived macrophages, VR1814_GF_ showed the strongest induction of cell-cell fusion in both M1 and M2 macrophages, followed by FIX-BPC and FIX-BP (Fig. 6C, D).

**Figure 6.**
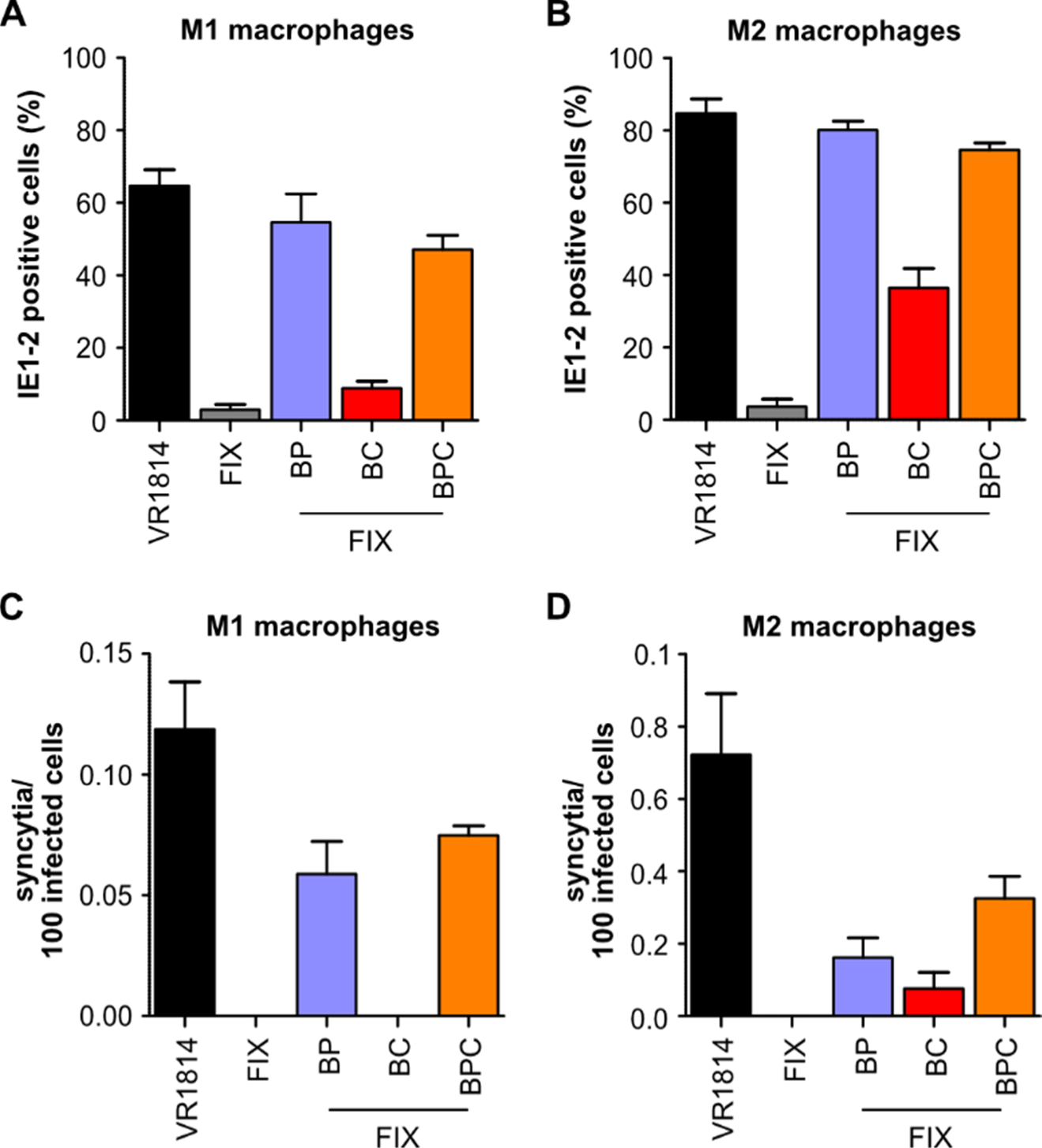
Infectivity of recombinant FIX strains and ability to induce syncytium formation in monocyte-derived macrophages. (A, B) M1 and M2 macrophages were infected at a MOI of 5. Relative infectivity of recombinant FIX strains was analyzed at 1 dpi in M1 and M2 macrophages, respectively, as percentages of IE1/IE2-positive nuclei. (C, D) Syncytium formation was quantified at 5 dpi by microscopic inspection and counting in M1 and M2 macrophages, respectively. Bars represent the mean ± SEM of four experiments with macrophages from different blood donors.

## Discussion

HCMV is a herpesvirus with remarkable natural variability (49, 50). Viral strains propagated in cell culture also show phenotypic differences, for example in their ability to infect and replicate in various cell types. These phenotypic differences can either be a consequence of natural variability or the result of selection and adaptive mutations in cell culture. It has long been known that the extensively passaged HCMV laboratory strains AD169 and Towne have accumulated numerous mutations that allow efficient virus replication in fibroblasts, but preclude infection of many other cell types (51). Mutations in RL13 and UL128L, which promote the release of cell-free virus, are primarily responsible for this property. Low-passage clinical strains, by contrast, are usually propagated in endothelial or epithelial cells in order to preserve the pentameric complex. Nevertheless, these strains may have undergone adaptive changes that favor virus replication in these cells (42, 45). For this reason, no cell culture-passaged virus is identical to the virus that was originally present in a human patient. There is also a broad consensus in the field that no HCMV strain can be considered a universal prototype, as no single strain can reflect the natural diversity of HCMV (23, 52). The most suitable viral strain for research purposes must be selected depending on the specific research question.

HCMV strain VR1814 was isolated in 1996 from the cervical secretions of a pregnant woman in Italy (28). The virus was passaged on endothelial cells to preserve its tropism for non-fibroblast cells. Indeed, VR1814 remained highly infectious for endothelial cells, epithelial cells, and cells of the myeloid lineage such as monocytes, macrophages, and dendritic cells. VR1814 was the first low-passage HCMV strain to be cloned as a BAC (32). Unfortunately, the properties of VR1814 were incompletely preserved in the VR1814-derived FIX-BAC: the virus retained only low-level infectivity for non-fibroblast cells and grew to rather modest titers in cell culture. Although FIX-BAC was initially used by many labs, its use decreased as BAC clones of other low-passage strains, such as TB40/E, TR, and Merlin (33–35), became available. A systematic genetic study has documented the genetic changes associated with the sequential passaging of VR1814 in endothelial cells and fibroblasts (42). However, the phenotypic consequences of the adaptive genetic changes have not been investigated. In the present study, we compared the phenotypes and genome sequences of the endothelial cell-passaged VR1814_GF_ and FIX. We demonstrated that a combination of two genetic loci, UL55/gB, and UL128L, of VR1814_GF_ confer high infectivity towards epithelial cells and macrophages when incorporated into FIX. The same two loci also confer the ability to form syncytia in epithelial cells and macrophages. One of the three amino acid exchanges in UL128L, the P72S substitution in UL130, was most important for increasing the infectivity of FIX as it facilitated the incorporation of the pentameric complex into viral particles (Fig. 4). Moreover, a frameshift mutation in US28, leading to a truncated US28 protein, also increased IE1/2 expression in macrophages (Figs. 5 and 6). Why this mutation increases IE gene expression is not known. However, US28 is known to play a crucial role in the establishment and maintenance of latency in myeloid cells (53, 54), and a previous study has shown that a C-terminally truncated US28(1–314) protein displays an altered signaling activity (43). Hence, it stands to reason that the truncated US28 protein promotes IE gene expression and the lytic cycle in monocyte-derived macrophages. It is worth noting that FIX-BP and FIX-BPC resemble VR1814_GF_ in terms of infectivity and ability to induce cell-cell fusion, but do not reach quite the same levels. This is also true for viral replication kinetics in ARPE-19 epithelial cells (Fig. 2). It seems likely that differences in other viral genes contribute to the phenotype of VR1814_GF_. Candidates are, for instance, the variants of gH, a component of the core fusion machinery (55), and UL116, a chaperone promoting the incorporation of gH/gL complexes into virions (56, 57).

To address the question of whether the gene variants of VR1814_GF_ conferring increased infectivity and fusogenicity are naturally occurring variants selected by passage on endothelial cells or adaptive mutations, we compared them to all HCMV sequences in GenBank. We did not find any other strain with a similar US28 truncation as the one in VR1814_GF_, suggesting that this is an adaptive mutation acquired during passaging in cell culture. Similarly, the S72P substitution in UL130 of FIX, which is responsible for the low infectivity of FIX, is not present in any other HCMV strain, nor is it present in any of the sequenced VR1814 substrains (42). Hence, it seems likely that this mutation has occurred during BAC cloning. However, the S585G substitution in gB and the T203N substitution in UL130 are not unique to VR1814_GF_ but are also present in three other HCMV strains (GenBank MT070138, MT070141, MT070142). At least two of these strains induce syncytium formation (58). Like VR1814, they were isolated in northern Italy, suggesting that these glycoprotein variants might exist as natural variants in a specific geographical region.

Ideally, virus research should be conducted with a strain that is as close as possible to the clinical pathogen. In this respect, HCMV research is in a dilemma (discussed previously in (23)). Fresh clinical HCMV isolates retain their wild-type properties for only a few passages (42) and are highly cell-associated. Therefore, they cannot be readily used in experimental research. Conversely, passaged virus strains produce higher titers of cell-free virus and are therefore more practical for experimental research, but their properties do not fully reflect the clinical pathogen. Since this dilemma cannot be avoided, a certain compromise is necessary to enable experiments (23). Passaged strains that grow to high titers and exhibit broad tropism, such as TB40/E and VR1814, are still needed for studies that require efficient infection of endothelial, epithelial, or myeloid cells (52). BAC clones of such strains are necessary to allow genetic modifications. The BAC clone of TB40/E, TB40-BAC4, is widely used in HCMV research. In contrast, the BAC clone of VR1814, FIX-BAC, has fallen out of favor due to its low infectivity for relevant cell types. The modified FIX-BACs described in this study, FIX-BP, and FIX-BPC, have much higher infectivity for epithelial and macrophages, similar to the parental VR1814. Hence, we propose that these “fixed” FIX strains should be particularly useful for infection studies on myeloid cells and for investigating the mechanisms and consequences of cell-cell fusion.

## Materials and Methods

### Cell Culture

Primary human foreskin fibroblasts (HFF-1, SCRC-1041), human embryonic lung fibroblasts (MRC-5, CCL-171), retinal pigmented epithelial cells (ARPE-19, CRL-2302), and human primary umbilical vein endothelial cells (HUVEC, PCS-100-010) were obtained from the American Type Culture Collection (ATCC). HFF cells were grown in Dulbecco’s modified Eagle’s medium (DMEM) (PAN-Biotech, Aidenbach, Germany), supplemented with 5% fetal calf serum (FCS) (Gibco, Waltham, MA, USA), 0.5 ng/mL of recombinant human fibroblast grow factor-basic (FGF-basic) (Peprotech, Rocky Hill, NJ, USA), 100 U/mL of penicillin, and 100 μg/mL of streptomycin. ARPE-19 cells were cultured in DMEM/F-12 GlutaMAX (Gibco) supplemented with 10% FCS, 100 U/mL penicillin, 100 μg/mL streptomycin, 15 mM HEPES, and 0.5 mM sodium pyruvate. HUVEC cells were cultured in cell culture flasks coated with gelatin in endothelial cell basal medium supplemented with 2% FCS and growth factors (Endothelial Cell Growth Medium kit) (PromoCell, Heidelberg, Germany). For infection experiments, cells were cultured in DMEM medium supplemented only with FCS and antibiotics.

The THP-1 monocytic cell line (ACC 16) was purchased from the German Collection of Microorganisms and Cell Cultures (DSMZ). THP-1 cells were propagated in RPMI 1640 GlutaMAX medium (Gibco) supplemented with 10% FCS, 100 U/mL penicillin, 100 μg/mL streptomycin, 10 mM HEPES, 1 mM sodium pyruvate, and 50 μM β-mercaptoethanol. For infection experiments, cell differentiation into macrophages was induced by adding 50 nM phorbol 12-myristate 13-acetate (PMA) (Sigma-Aldrich, St. Louis, MO, USA) to the medium for 72 hours.

Buffy coats of healthy HCMV-seronegative blood donors were obtained from the Transfusion Center of the University Medical Center Hamburg-Eppendorf (Hamburg, Germany). M1 and M2 macrophages were generated as previously described (59). Briefly, human monocytes were isolated by negative selection with magnetic microbeads according to the manufacturer’s instructions (Pan Monocyte Isolation Kit) (Miltenyi Biotec, Bergisch Gladbach, Germany). A total of 3 x 10^6^ monocytes/mL were seeded in complete RPMI medium in hydrophobic Lumox dishes (Sarstedt, Nümbrecht, Germany), and differentiated into M1 or M2 macrophages by incubation for 7 days in the presence of 100 ng/mL recombinant human granulocyte-macrophage colony-stimulating factor (GM-CSF) or macrophage-colony stimulating factor (M-CSF) (R&D Systems, Minneapolis, MN, USA), respectively.

### Viruses and stock preparation

The HCMV strains TB40/E (60) and VR1814 (28) were obtained from Christian Sinzger (University of Ulm, Germany) and Giuseppe Gerna (Policlinico San Matteo, Pavia, Italy). The VR1814 variant used in our lab was named VR1814_GF_. FIX-GFP is a modified FIX-BAC (32) containing a GFP expression cassette within the BAC vector backbone (38). For the preparation of cell-free virus stocks, HFF cells were co-cultured with late-stage infected HUVEC cells at a ratio of roughly 50:1. Supernatants were harvested from infected cultures showing 100% cytopathic effect. Cellular debris was removed by centrifugation at 5500 × *g* for 15 min, virus particles were pelleted by ultracentrifugation at 23,000 × *g* for 60 min at 4°C. Virus pellets were resuspended in DMEM medium containing a cryopreserving sucrose-phosphate buffer (74.62 g/L sucrose, 1.218 g/L K_2_HPO_4_, 0.52 g/L KH_2_PO_4_), aliquoted, and stored at −80°C. Viral titers were determined as previously described (61). Briefly, HFF or ARPE-19 cells were inoculated with serial dilutions of the virus stocks. After 48 hours, the cells were fixed with methanol/acetone for 10 min at −20°C, blocked with PBS/1% milk for 15 min, and incubated with an anti-IE1/2 antibody for 2 hours at 37°C and an HRP-conjugated rabbit anti-mouse secondary antibody (Jackson ImmunoResearch, Cambridge, UK) for 45 min at 37°C. Staining was done with 3-amino-9-ethylcarbazole (AEC) (BioLegend, Amsterdam, The Netherlands) as HRP substrate. IE1/2 positive nuclei were counted, and viral titers were determined as infectious units per mL (IU/mL).

### Plasmids and Reagents

The cloning vector pBR322 was a gift from Adam Grundhoff (Leibniz Institute of Virology, Hamburg, Germany). pEPkan-S and *E. coli* strain GS1783 have been described previously (62). GS1783 bacteria were grown in LB broth (Lennox) (Carl Roth, Karlsruhe, Germany) containing 5 g/L of NaCl (Sigma-Aldrich). Lentiviral vector plasmids encoding DSP1–7 or DSP8–11 and their helper plasmids pMD-G and pCMVR8.91 were kindly provided by Dalan Bailey (The Pirbright Institute, Woking, UK). Antibiotics were purchased from Roth or Invitrogen and used at the following concentrations: ampicillin (100 µg/mL), kanamycin (50 µg/mL), chloramphenicol (15 µg/mL), and zeocin (25 µg/mL). L-(+)-arabinose was purchased from Sigma-Aldrich.

### Viral genome sequencing

Viral genome sequences were determined by Illumina sequencing at the Next Generation Sequencing facility of the Leibniz Institute of Virology essentially as described previously (63).

### Mutagenesis of HCMV genomes

Point mutations of specific regions of interest were introduced into the BAC clone of VR1814, FIX, by *en passant* mutagenesis (62). The UL128 locus of VR1814 was cloned and used to replace UL128L in FIX, essentially as described (64). All modified FIX strains were examined by restriction fragment analysis and sequencing of the mutated region. FIX-derived viruses were then reconstituted in fibroblasts and propagated in HUVEC cells. The complete genome sequences of VR1814 (accession no. GU179289) and FIX-BAC (accession no. AC146907) are available at GenBank.

### Replication kinetics

One day prior to infection, ARPE-19 cells were seeded in 6-well plates at a density of 1.2 × 10^5^ cells/well. Cells were infected in triplicates at an MOI of 0.5 for 3 hours at 37°C, washed with phosphate-buffered saline (PBS) (Sigma-Aldrich), and supplied with fresh medium. Supernatants were harvested at indicated time points and titrated on HFF cells as described above.

### Immunodetection

The following monoclonal antibodies were used in this study: anti-IE1/2 (3H4) and anti-pp65 (8F5) were provided by Thomas Shenk (Princeton University, Princeton, NJ, USA). Anti-pp150 (XPA 36-14) was provided by William Britt (University of Alabama, Birmingham, AL, USA). Anti-UL128 (4B10) and anti-UL130 (3E3) antibodies (44) were provided by Barbara Adler (University of Munich, Munich, Germany). Antibodies against UL44 (10D8) and GAPDH (14C10) were purchased from Virusys (Milford, MA, USA) and Cell Signaling (Beverly, MA, USA), respectively.

Immunoblotting was performed according to standard protocols. At different times post-infection, cells were lysed with RIPA buffer (50 mM of Tris-HCl, pH 8; 150 mM of NaCl, 1mM EDTA, 1% NP-40, 0.5% deoxycholate, 0.1% SDS) supplemented with cOmplete Mini Protease Inhibitor Cocktail (Roche, Penzberg, Germany). Cell-free virions were further cleared of debris by centrifugation at 1000 × *g* for 5 min at 4°C and resuspended in RIPA buffer supplemented with cOmplete Mini Protease Inhibitor Cocktail. Insoluble material from all lysate samples was removed by centrifugation at 16,000 × *g* for 10 min, and the cleared extracts were heated to 95°C for 5 min. For reducing blots, extracts were supplemented with 1% β-mercaptoethanol. Equal protein amounts of the different samples were separated by sodium dodecyl sulfate-polyacrylamide gel electrophoresis (SDS-PAGE) and transferred onto nitrocellulose or polyvinylidene difluoride (PVDF) membranes (Bio-Rad Laboratories, Feldkirchen, Germany) by semidry blotting. Proteins of interest were detected with specific primary antibodies and secondary antibodies coupled to horseradish peroxidase (Jackson ImmunoResearch). Densitometry was measured with ImageJ (https://imagej.net).

Immunofluorescence analysis was performed as previously described (40). At different times post-infection, cells were fixed with methanol/acetone, blocked with 1% milk in PBS, and incubated with specific primary antibodies overnight at 4°C or for 2 h at 37°C, and secondary antibodies coupled to AlexaFluor 555 or 647 (Thermo Fisher Scientific, Schwerte, Germany). DAPI (4’,6-diamidino-2-phenylindole) (Sigma-Aldrich) was used to stain nuclear DNA. Images were acquired using a Nikon A1+ LSM confocal microscope. For investigating the infectivity, fluorescence images were acquired using a CellInsight CX5 High-Content Screening Platform (Thermo Fisher Scientific). The percentage of IE1/2-positive cells was determined by using HCS Studio software.

### Quantification of HCMV genomes

Cell-free HCMV stocks (50 µL) were first treated with turbo DNase (Thermo Fisher Scientific) to remove DNA not contained within viral particles. Total DNA was extracted using an InnuPREP DNA Mini Kit (Analytik Jena, Jena, Germany), and eluted in 50 µL of nuclease-free water. Viral genome copies were quantified by real-time quantitative PCR (qPCR) using SYBR green dye (Thermo Fisher Scientific) and primers specific for the HCMV UL36 ORF (ACGCAAAGAGTTCCTCGTAC and TGAACATAACCACGTCCTCG). PCR products were detected using a QuantStudio 3 (Thermo Fisher Scientific) qPCR machine. Serial dilutions of a FIX-BAC DNA were used as a reference to calculate the genome copies/mL as described (39). For each experiment, two independent viral DNA extractions and three independent qPCRs were performed.

### Cell-cell fusion assay

ARPE-19 cells expressing the dual split protein (DSP) system were generated by transduction with lentiviral vectors essentially as described previously (40). HEK293T cells were transfected with lentiviral vector plasmid, pCMVR8.91, and pMD-G. Lentivirus-containing supernatants were used to transduce ARPE-19 cells. Transduced cells were selected with 1 µg/mL of puromycin (Sigma-Aldrich).

For the fusion assay, equal numbers of ARPE-19 cells expressing DSP1-7 or DSP8-11 were combined and infected with HCMV at an MOI of 1. At 5 dpi, cells were washed with PBS, and incubated with coelenterazine-h (Promega, Madison, WI, USA) at a final concentration of 2.5 nM. To quantify *Renilla* luciferase activity, luminescence was measured on a multi-mode microplate reader (FLUOstar Omega) (BMG Labtech, Ortenberg, Germany).

### Syncytium formation inhibition assay

One day prior to infection, DSP-expressing ARPE-19 cells were seeded in 96-well plates at a density of 1 × 10^4^ cells/well. Cells were infected with HCMV at an MOI of 1 for 3 hours at 37°C, washed with PBS, and incubated with fresh medium containing serial dilutions of anti-UL130 antibody (3E3 hybridoma supernatant). At 5 dpi, *Renilla* luciferase expression was quantified as described above, and the percentage of syncytia relative to untreated cells was calculated. Cells were then fixed and stained with an anti-IE1/2 antibody and DAPI. Fluorescence images were acquired using CellInsight CX5 High-Content Screening Platform.

## Acknowledgments

We thank Thomas Shenk, William Britt, and Barbara Adler for antibodies, Dalan Baley for lentiviral vector plasmids, and Roland Thünauer and Marcel Schie (LIV Microscopy and Image Analysis facility) for their help and support. Schematic figures were created with BioRender. This research was supported by the Deutsche Forschungsgemeinschaft (grant BR1730/9-1 to W.B.). G.C. was supported by a Leibniz Center Infection (LCI) scholarship.

Conceptualization, G.C., W.B., and G.F.; methodology, G.C., X.Z., and G.F.; investigation, G.C., X.Z., and G.F.; writing - original draft preparation, G.C., and W.B.; writing - review and editing, all authors; supervision, W.B. and G.F.; funding acquisition, W.B. and G.F.

